# Exploring the accuracy and limits of algorithms for localizing recombination breakpoints

**DOI:** 10.1101/2023.12.08.570844

**Authors:** Shi Cen, David A. Rasmussen

## Abstract

Phylogenetic methods are widely used to reconstruct the evolutionary relationships among species and individuals. However, recombination can obscure ancestral relationships as individuals may inherit different regions of their genome from different ancestors. It is therefore often necessary to detect recombination events, locate recombination breakpoints and select recombination-free alignments prior to reconstructing phylogenetic trees. While many earlier studies examined the power of different methods to detect recombination, very few have examined the ability of these methods to accurately locate recombination breakpoints. In this study, we simulated genome sequences based on ancestral recombination graphs and explored the accuracy of three popular recombination detection methods: MaxChi, 3SEQ and GARD. The accuracy of inferred breakpoint locations was evaluated along with the key factors contributing to variation in accuracy across data sets. While many different genomic features contribute to the variation in performance across methods, the number of informative sites consistent with the pattern of inheritance between parent and recombinant child sequences always has the greatest contribution to accuracy. While partitioning sequence alignments based on identified recombination breakpoints can greatly decrease phylogenetic error, the quality of phylogenetic reconstructions depends very little on how breakpoints are chosen to partition the alignment. Our work sheds light on how different features of recombinant genomes affect the performance of recombination detection methods and suggests best practices for reconstructing phylogenies based on recombination-free alignments.

## Introduction

Recombination exchanges genetic material horizontally between organisms and thus plays a crucial evolutionary role in creating novel genotypes (Posada *et al*., 2002). Because recombination allows for individuals to inherit different regions of their genomes from different parents, recombination can bring beneficial alleles from different genomic backgrounds together, increasing the rate of adaptation (Alves *et al*., 2017). Recombination also allows for deleterious mutations to be purged, avoiding the accumulation of deleterious mutations in a single genetic background known as Muller’s ratchet (Muller, 1964). Detecting the occurrence of recombination is therefore important to understanding the evolutionary history of genomic sequence data and estimating population genetic parameters such as the recombination rate which fundamentally shape the trajectory of evolving populations (Visscher and Hill, 2006).

Recombination can also mislead the inference of phylogenetic relationships from sequence data. In the presence of recombination, the true ancestral relationships among individuals may vary across the genome because individuals inherit different regions of their genome from different ancestors. No single phylogeny can therefore represent the true ancestral relationships among individuals across a recombining genome (Awadalla, 2003; Rasmussen *et al*., 2014). If recombination is ignored during phylogenetic reconstruction, the inferred phylogeny may not accurately reflect the true relationships for any region of the genome (Kubatko and Degnan, 2007). Moreover, ignoring recombination during phylogenetic reconstruction can distort the topology of inferred phylogenetic trees, leading to more star-like trees with longer terminal branches and less clock-like rates of molecular evolution (Schierup and Hein, 2000). Hence, it is essential to detect recombination events prior to performing phylogenetic analysis and then either remove recombinant sequences entirely or partition sequence alignments into recombination-free segments (Bell and Bedford, 2017; Boni *et al*., 2020; Jackson *et al*., 2021).

Because of the central role of recombination in biological sequence analysis, several algorithms have been developed to detect recombination events. These algorithms can be divided into a few major classes depending on what information and statistical methods are used to detect recombination (Shikov *et al*., 2022; Posada and Crandall, 2001). These include: (1) Substitution distribution methods which aim to detect a mosaic structure in the distribution of substitutions (i.e. polymorphic sites) across the genome. For example, MaxChi uses a *χ*^2^ statistic to test whether there are significantly more substitutions between pairs of sequences to the left or right of a potential breakpoint than would be expected by chance (Smith, 1992). Similar ideas can be applied to sequence triplets. For example, 3SEQ considers whether substitutions between a presumed child sequence and two parent sequences are clustered such that the child sequence resembles different parental sequences over the span of the genome (Boni *et al*., 2007). (2) Phylogenetic methods aim to identify recombination breakpoints by looking for phylogenetic discordance between trees reconstructed from different regions of a genome. For example, GARD iteratively divides a genome into smaller blocks, builds neighbor-joining trees for each block, and then tests whether the sequence data strongly supports discordance between trees indicative of possible recombination (Pond *et al*., 2006). Lastly one can consider “compatibillity” methods which consider whether each site in a sequence is congruent with an evolutionary history without recombination and recurrent mutations (Hudson and Kaplan, 1985; Smith and Smith, 1998). However, since recurrent mutations, including convergence and reversion, are a frequent cause of homoplasy in rapidly evolving populations, we do not consider compatibility methods here.

Regardless of the exact method employed, detecting and precisely locating recombination breakpoints is challenging. Only a small fraction of sites in a given sequence alignment are likely to be informative about recombination. At a minimum, there needs to be polymorphic sites present in two recombining parental sequences in order for a signal of recombination to be detected since it is not possible to detect recombination among identical sequences. Furthermore, not all polymorphic sites will necessarily be informative about recombination because processes such as recurrent mutation may obscure signals of recombination. There has therefore been great interest in evaluating the performance of recombination detection methods by analyzing their detection power (sensitivity) and false positive rate (specificity) on real and simulated sequences (Wiuf *et al*., 2001; Posada and Crandall, 2001; Posada, 2002; Bay and Bielawski, 2011). All these studies have shown that detection methods can under certain circumstances (e.g. high sequence diversity) perform well in terms of power and false positive rates. However, these previous studies paid little attention to how accurately and precisely different methods could localize recombination breakpoints. This is an important oversight as the ability to localize breakpoints has many practical applications in bioinformatics including creating non-recombinant alignments for phylogenetic reconstruction.

When recombination detection methods are used to identify breakpoints prior to phylogenetic reconstruction, the location of identified breakpoints are generally used to partition or “slice” the full alignment into local sub-alignments presumed to be free of recombination, which are then used to reconstruct local phylogenetic trees for non-recombinant regions of the genome (Tamura *et al*., 2023). However, one can choose between several different strategies for slicing alignments based on identified breakpoints. For example, one could be as aggressive as possible and slice the alignments at the exact position of identified breakpoints in order to include as many sites as possible in the resulting sub-alignments. Alternatively, one could be as conservative as possible and only select sub-alignments that are free of recombination with high-confidence, excluding all sites within a certain confidence window around identified breakpoints. To our knowledge, how errors in breakpoint localization impact downstream phylogenetic inference and how to best choose recombination-free subalignments based on identified breakpoints has never been systematically explored.

In this paper, we use simulated sequence data to explore the accuracy and precision of recombination detection methods including MaxChi, 3SEQ and GARD at localizing recombination breakpoints. Our simulation study consists of three parts. In the first part, we simulate triplets of sequences with only a single recombination breakpoint under the coalescent with recombination (Hudson, 1983; Hudson *et al*., 1990). We then investigate which genomic features lead to the greatest variability in performance/accuracy between different data sets and methods. In the second part, we consider performance among the same methods in the more general setting where multiple recombination events generate the observed sequence data. In the third part, we explore how best to partition alignments into recombination-free segments based on the breakpoints identified by different detection methods. Through these three simulation studies, we aim to provide insights into the selection of appropriate recombination detection methods and best practices for subsequently reconstructing phylogenetic trees based on inferred breakpoint locations.

## Related Work

We briefly summarize how each of the three recombination detection algorithms explored here test for the presence of recombination and localize the most likely recombination breakpoints.

The MaxChi method, developed by Smith (1992), is a classic substitution distribution method. For a single recombination event, it assumes that the recombinant genome consists of two blocks inherited from two independent parent sequences. For a sequence pair, it makes an arbitrary cut at position *k*, and builds a 2 × 2 contingency table around the position *k*, to find the statistic *χ*^2^ that indicates whether the number of mutations to the left/right of the breakpoint deviates from what would be expected based on the total number of mutations in each region under the null hypothesis that there is no recombination events. The site *k_max_* that maximizes the *χ*^2^ statistic is considered a candidate breakpoint location. If there are multiple breakpoints to infer, MaxChi will iteratively search each block until no further breakpoints are detected.

Genetic Algorithm Recombination Detection (GARD) is a classic phylogenetic-based recombination detection method developed by Pond *et al*. (2006). At each site in the alignment, GARD divides the alignment into two blocks and builds a neighbor-joining (NJ) tree individually for each block. GARD then uses corrected Akaike information criterion (*AIC_c_*) values to decide if the likelihood of the sequence data given two trees is significantly higher than under a single tree. If at least one *AIC_c_* value of the inferred NJ trees from two blocks is less than the *AIC_c_* value of the NJ tree inferred from the whole alignment, that position will be considered a potential breakpoint. When dealing with multiple breakpoints, GARD uses a genetic algorithm to search for candidate positions based on the likelihood of the model (Pond and Frost, 2005; Eshelman, 1991).

3SEQ is a widely used exact nonparametric method for detecting recombination developed by Boni *et al*. (2007). Given three sequences, one sequence (C) is presumed to be the recombinant child while the other two sequences, denoted as P and Q, are presumed to be the parents of C. 3SEQ considers the sequence of mutations in C that match the allelic state of the parents P or Q at a given site. A sequence of several P-like mutations followed by a sequence of several Q-like mutations would therefore suggest C inherited genetic material from P to the left and from Q to the right of a recombination breakpoint. Under the null hypothesis that there is no recombination, the algorithm calculates the exact p-value of observing a sequence of P or Q-like mutations in the child sequence based on a hypergeometric random walk model. A very unlikely sequence of mutations under the null hypothesis of no recombination (low p-values) is considered to be evidence for a recombination event. The test is then repeated for all possible sequence triplets in an alignment.

## Methods

### Ancestral Recombination Graph Simulation and Sequence Simulation

To simulate sequence alignments, we used msprime (Baumdicker *et al*., 2021) to first simulate ancestral recombination graphs (ARGs) with known recombination events. In our first set of simulations, we fixed the sample size at three and conditioned the simulations on having exactly one recombination event occur between two parent lineages and a child recombinant sequence. The location of the true recombination breakpoint was recorded for comparison with the inferred locations. After simulating each ARG, the pyvolve package (Spielman and Wilke, 2015) was used to simulate sampled sequences along each local tree in a given ARG and then sequences were concatenated end-to-end to obtain a full alignment spanning the entire genome. We varied the mutation rate *µ* such that pairwise genetic diversity *π* ranged from 0.01 to 0.2. All other simulation parameters are given in Table 1. In our second set of simulations we constrained the number of recombination events between 1 and 10 such that there could be multiple breakpoints in the simulated alignments. For these simulations, we expanded the pairwise genetic diversity range up to 0.3 and set the sample size to 10. Simulated ARGs that contained undetectable recombination events (i.e. the Type I events defined below) were discarded. All simulation code and data is available at https://github.com/davidrasm/RecBreakDetect.

**Table 1:**
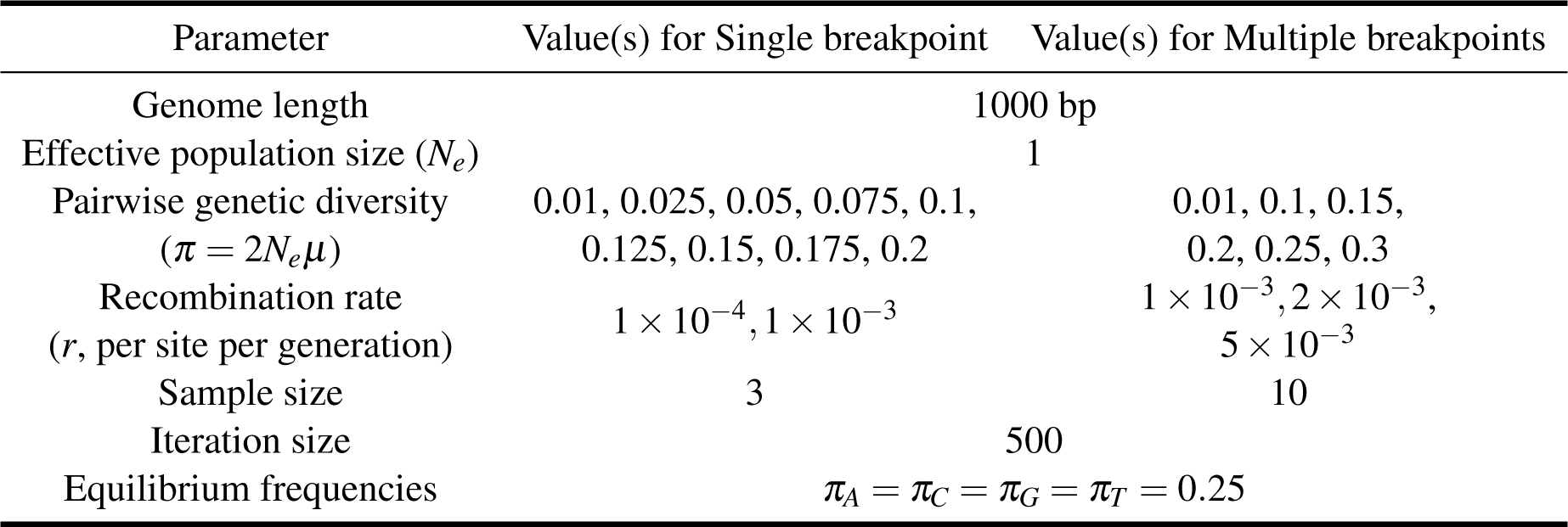
Parameters used to simulate ARGs and sequence alignments.

| Parameter | Value(s) for Single breakpoint | Value(s) for Multiple breakpoints |
| --- | --- | --- |
| Genome length | 1000 bp |  |
| Effective population size ($N_e$) | 1 | |
| Pairwise genetic diversity ($\pi = 2N_e\mu$) | 0.01, 0.025, 0.05, 0.075, 0.1, 0.125, 0.15, 0.175, 0.2 | 0.01, 0.1, 0.15, 0.2, 0.25, 0.3 |
| Recombination rate ($r$, per site per generation) | $1 \times 10^{-4}$, $1 \times 10^{-3}$ | $1 \times 10^{-3}$, $2 \times 10^{-3}$, $5 \times 10^{-3}$ |
| Sample size | 3 | 10 |
| Iteration size | 500 |  |
| Equilibrium frequencies | $\pi_A = \pi_C = \pi_G = \pi_T = 0.25$ | |

### Recombination Breakpoint Detection and Performance Evaluation Metrics

Three methods were applied to detect recombination events: MaxChi (Smith, 1992), 3SEQ (Boni *et al*., 2007) and GARD (Pond *et al*., 2006). Simulations in which the recombination event was not identified by any method were discarded. We then used two evaluation metrics to compare their relative performance in localizing breakpoints. First, we define the error as the distance of the imputed breakpoint location from the true location in units of base pairs. For single breakpoint simulations, we measure the localization error as the distance between the true breakpoint and the inferred breakpoint from three recombination detection methods. For multiple breakpoint simulations, different measurement strategies are used depending on the detection method. For 3SEQ, each row in a 3SEQ output file indicates a potential recombination event between three sequences. Within the full simulated ARG, we identify the sub-ARG that only contains these three sequences and check whether there is a recombination event in this sub-ARG. If so, for each inferred breakpoint interval, we calculate the distance between the midpoint of the interval and the nearest recombination breakpoint in the sub-ARG. An average localization error for each simulation is calculated from all of the intervals. A similar strategy is used for MaxChi: for all pairs of sequences in which a recombination event is detected we look at the corresponding sub-ARG involving the two sequences and find the nearest true recombination breakpoint to calculate the mean localization error. For GARD, we directly find the nearest recombination breakpoint for each inferred breakpoint to calculate the mean localization error.

Second, we define precision as the reciprocal of the variance in the site-specific probabilities of each site being the recombination breakpoint, such that placing all probability on a single site would have the highest precision and placing uniform probability on all sites would have the lowest precision. The site-specific probabilities were normalized such that these probabilities sum to one. How site-specific breakpoint probabilities were computed for each method is described in the Supplemental material 1.

### Explaining variability in performance across data sets

We looked at several summary statistics or features of the simulated sequence data that could potentially explain the variable performance of different methods across simulations. These features were used in the partial least squares regression model described below. These variables include:

#### Recombination type categorization

Following Wiuf and Hein (1999), we categorize recombination events into three types. An example ARG corresponding to each type is shown in Figure 1. Type I recombination events happen when the two parent lineages of a recombinant child immediately coalesce before they merge with other sampled lineages in the genealogy. In this case, recombination has no impact on local tree topologies and leaves no trace in the sequence data. Type I events should therefore be undetectable by any method, but we include them here to test the specificity of each method against detected false positives. Type II events happen when one of the parent lineages coalesces with another sampled lineages to form a new parent lineage that then immediately coalesces with the other parent lineage. Hence, there is no topology change on the local trees, but the branch lengths of the parent lineages may differ. Type III events happen when the recombinant lineage coalesces with different non-recombinant lineages on the two sides of the recombination breakpoint. This type results in a clear topology change between the two local trees.

**Figure 1:**
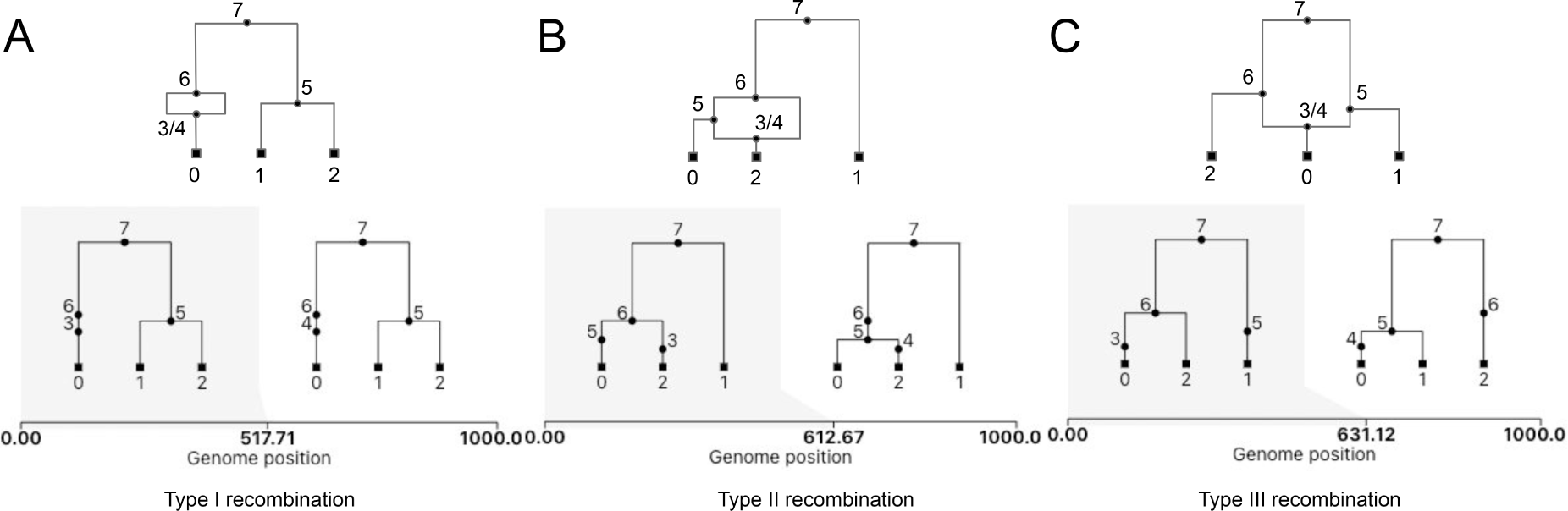
Illustration of the three recombination event types as categorized by Wiuf and Hein (1999). A: Type I recombination; B: Type II recombination; C: Type III recombination.

#### Number of consistent and inconsistent informative sites

For any sequence triplet, let the true recombinant child be denoted as *C*, the parental sequence inherited along the left side of the recombinant child sequence be designated as the *P* sequence and the parental sequence inherited along the right side be the *Q* sequence. We define informative sites as positions where *C* and either *P* or *Q* share the same mutation/nucleotide, but not both (Boni *et al*., 2007). We further define consistent informative sites as positions in *C* such that the nucleotide state of *C* matches the parent from which it inherited its genetic material at that site. Figure 2 illustrates our definition of consistent versus inconsistent informative sites. However, recurrent mutations and back mutations (reversions) can obscure the pattern of inheritance by causing the nucleotide state of the child to match another sequence from which it did not inherit its genetic material as shown in Figure 2. We will refer to these sites as inconsistent informative sites.

**Figure 2:**
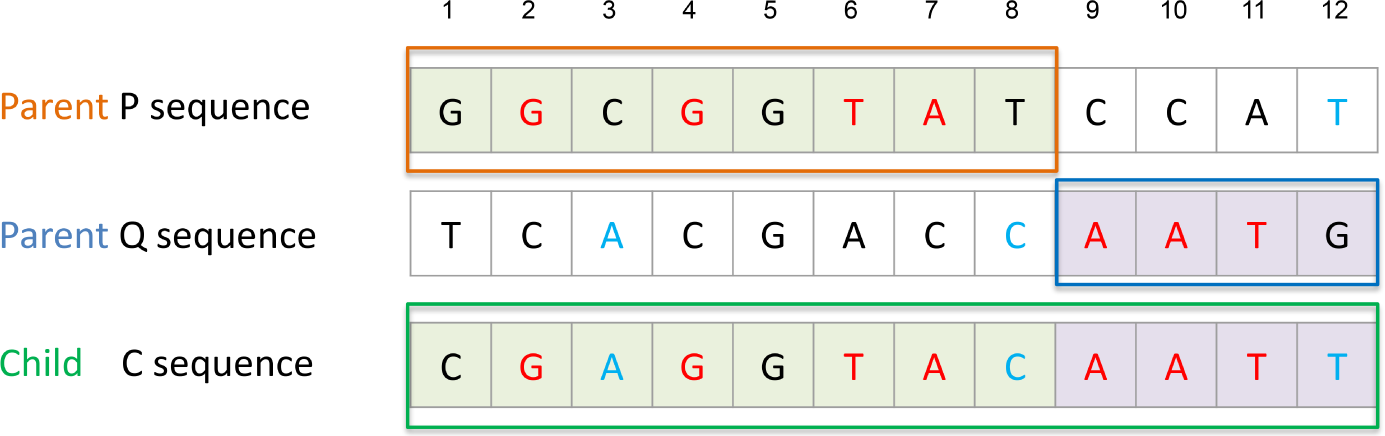
A hypothetical alignment resulting from a single recombination event between the parents P and Q producing the recombinant child sequence C. The consistent informative sites are colored red and the inconsistent sites are colored blue. For example, the eighth position is an informative site where the child has the same state as the *Q* parent. But the left side of the child sequence is actually inherited from the *P* parent. Hence, this site is considered to be an inconsistent informative site.

#### Average distance between informative sites and breakpoint

In order to measure the effect of the distance between informative sites and the true breakpoint location on detection performance, we define the average distance of informative sites as the mean value of the distance between each informative site and the true breakpoint.

### Partial Least Square Regression Model Building

We use a partial least squares regression (PLSR) model to evaluate the contribution of the 5 predictor variables described above to the performance of the recombination detection methods at localizing breakpoints. A PLSR model was chosen because there is strong multicollinearity among our predictor variables, a condition where standard linear regression often fails, but partial least squares excels because it seeks to find a linear combination of predictor variables that explains the maximum variance in the response variable (i.e. localization error) (Haenlein and Kaplan, 2004). As in Principle Component Analysis, the contribution of each explanatory variable to the new axes is quantified by a weight. Following the approach described in Ratmann *et al*. (2016), the explained variance of each predictor is calculated as 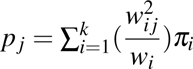, where *k* is the number of latent factors in each PLSR model for the three detection methods (*k* = 5 for GARD, MaxChi and 3SEQ via cross-validation), 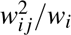 is the relative weight of the *j*th predictor in the *i*th latent factor with 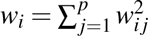 and *w_i j_* being the loading of the linear combination for each component, and *π_i_* is the explained variance by *i*th latent factor. We used the plsRglm package to fit the PLSR model and construct the confidence intervals of each coefficient using *R* = 5000 bootstraps (Bertrand and Maumy, 2023).

### Breakpoint detection for phylogenetic reconstruction

In order to evaluate how different partitioning strategies impact phylogenetic reconstruction, we performed simulation experiments in which we set different window sizes around the breakpoints identified by different detection methods. We then sliced the sequence alignment into local sub-alignments at either the true recombination breakpoint, the different window borders, or the midpoint of each window (Figure S2). Maximum likelihood trees were then reconstructed from the resulting local sub-alignments using RAxML (Stamatakis, 2014). The performance of the phylogenetic reconstruction was then evaluated by calculating the Robinson-Foulds (RF) distance (Robinson and Foulds, 1981) between the corresponding true local tree and the reconstructed local tree. For genomic regions inside windows excluded from any local sub-alignment for which a local tree could not be reconstructed, we use the neighboring ML trees to interpolate between regions in order to calculate the RF distance. All RF distances are normalized and averaged over each site. RF distances are calculated using the *symmetric_difference* function in DendroPy (Sukumaran and Holder, 2010).

## Results

### Detecting and localizing single breakpoints

#### An Overview of the Three Recombination Detection Methods

While previous studies explored the power of different algorithms to detect recombination events (Wiuf *et al*., 2001; Posada and Crandall, 2001; Posada, 2002), we begin our exploratory analysis by summarizing the power and specificity of GARD, MaxChi and 3SEQ. Because type I recombination events should be undetectable, we consider detection of type I events to be false positives. Failure to detect type II or III events are considered to be false negatives. Overall, MaxChi and 3SEQ show a high specificity (i.e. a lower false positive rate) at all levels of genetic diversity, whereas the specificity of GARD decreases as genetic diversity increases (Figure 3A). On the other hand, GARD shows a much greater power to detect recombination compared to the other two methods (Figure 3B). For type II events, MaxChi has a higher power than 3SEQ, while 3SEQ has a higher power than MaxChi when detecting type III events. We conclude that among the three detection methods, GARD is more sensitive at detecting recombination events, but there is a tradeoff between sensitivity and specificity, i.e. the increased sensitivity of GARD comes at the cost of a higher false positive rate.

**Figure 3:**
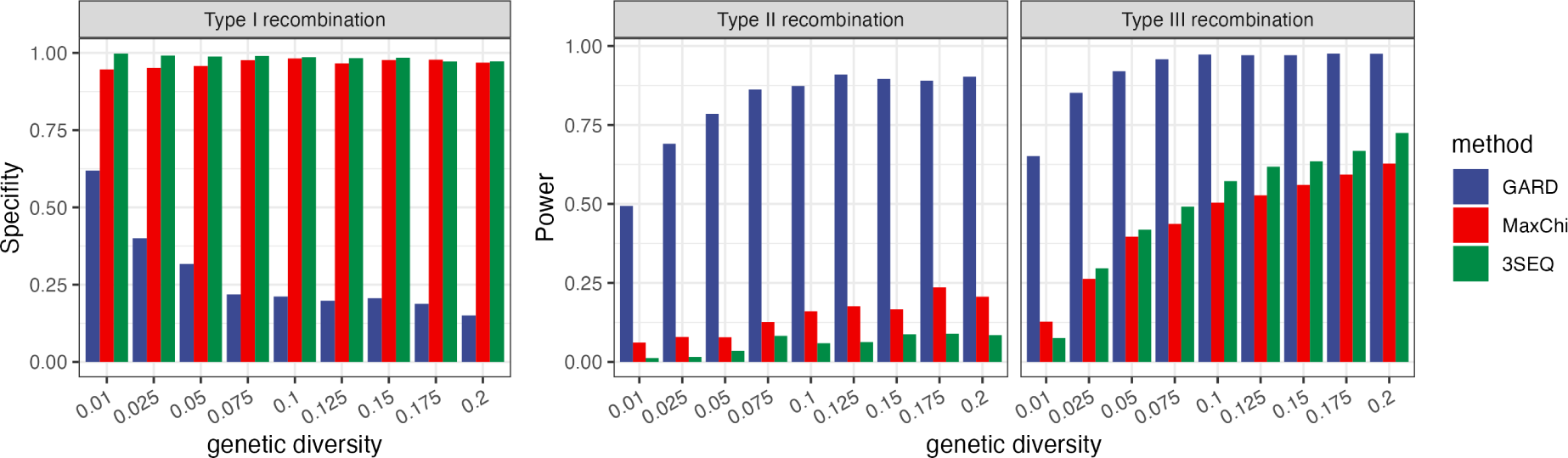
Sensitivity (power) and specificity of GARD, MaxChi and 3SEQ at detecting different types of recombination events at different levels of genetic diversity.

#### Accuracy in localizing breakpoints

To determine the accuracy of each method in localizing breakpoints we quantify the error in the inferred breakpoint location in terms of the distance in basepairs (bp) from the true location. The accuracy of the three detection methods in localizing recombination breakpoints substantially differs (Table 2). When detecting type I recombination events, MaxChi and GARD are no better than the expected error of a random guess, which should be approximately 333 bp given the probability of a breakpoint is uniformly distributed along a 1000bp sequence. For type II and III events, 3SEQ generally has a lower localization error (33.69bp) than the other two methods whereas GARD has a localization error of nearly 200bp.

**Table 2:** Localization accuracy (mean*±*SEM) of three detection methods averaging over all simulations at all levels of genetic diversity.

|  | Type I | Type II & Type III |
| --- | --- | --- |
| GARD | $364.66 \pm 4.47$ | $195.72 \pm 4.01$ |
| MaxChi | $395.27 \pm 22.40$ | $72.41 \pm 4.12$ |
| 3SEQ | $203.40 \pm 19.19$ | $33.69 \pm 2.07$ |

In order to see which features of the sequence data explain the most error in localizing breakpoints, we take the loading on each predictor variable in the PSLR model as a weight to evaluate the explained variance. Overall, 30% to 40% of the variance in performance can be captured by the features in the model for each of the three detection methods (Figure 4). How each feature contributes to localization error is summarized in Table 3 and explored individually below.

**Figure 4:**
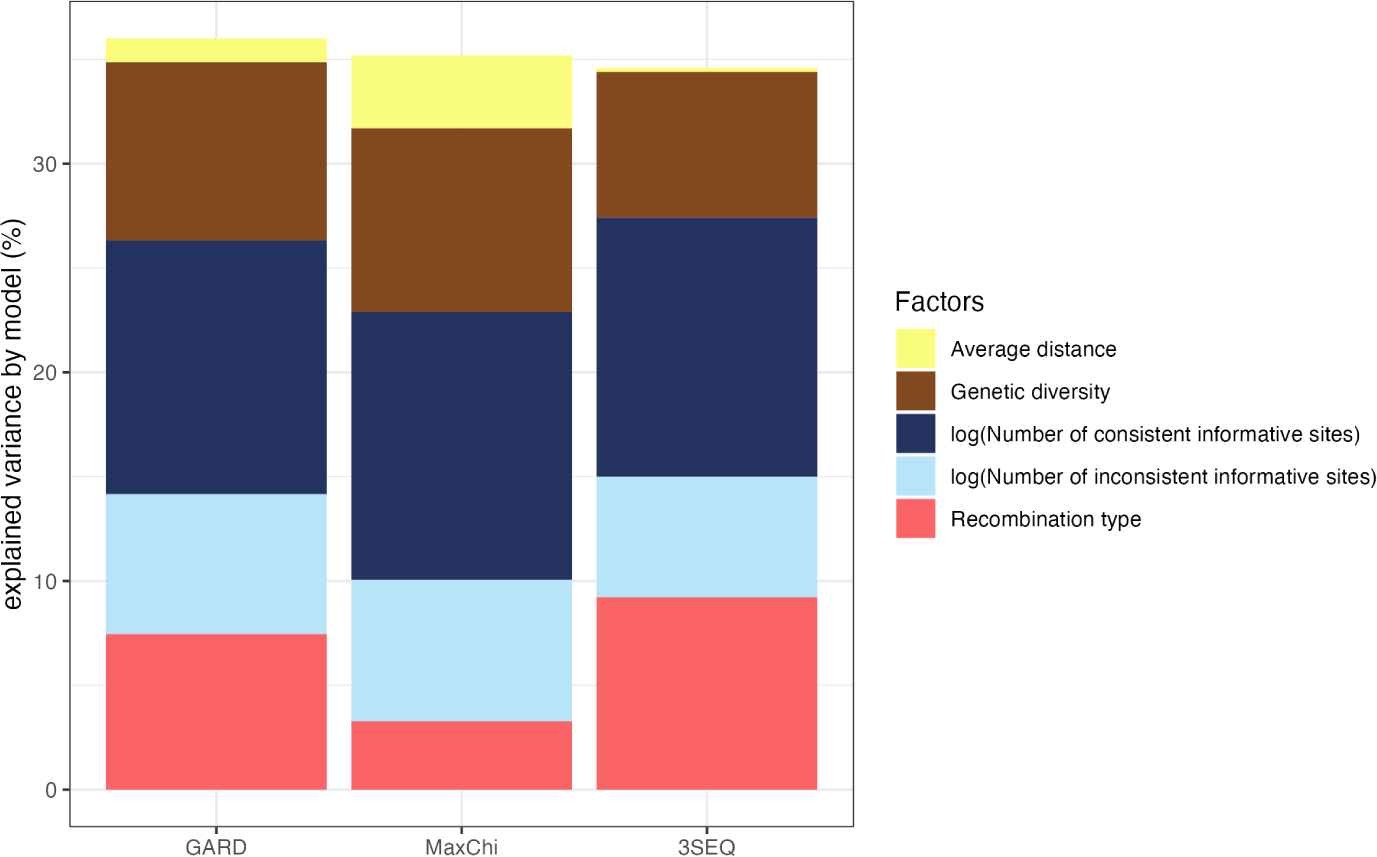
Variance in breakpoint localization error explained by each predictor variable for three detection methods as quantified by the PLSR model.

**Table 3:** Impact of each predictor on breakpoint localizaton error. The sign of the impact corresponds to the sign of the regression coefficient estimated under the PLSR model. Type II events are set as the reference level to type III events. Asterisks indicate significant effects.

| Predictor | GARD | MaxChi | 3SEQ |
| --- | --- | --- | --- |
| Genetic diversity | + | + | - |
| Recombination type | -* | -* | -* |
| Average distance between informative sites and breakpoint | +* | +* | - |
| Logarithm of number of consistent informative sites | -* | -* | -* |
| Logarithm of number of inconsistent informative sites | +* | +* | +* |

##### Recombination type

For all methods, error is higher for type II than type III events. This result makes intuitive sense as type II events only impact branch lengths and are therefore expected to leave a weaker signal in the sequence data than type III events which impact local tree topologies. Overall, MaxChi performs better when detecting type II events while 3SEQ has a significantly lower error when detecting type III events (ANOVA *P* < 0.001, Figure 5A). The *recombination type* factor explains more variance in performance for 3SEQ and GARD than for MaxChi (Table 4), as both 3SEQ and GARD are sensitive to the topology change in between two local trees.

**Figure 5:**
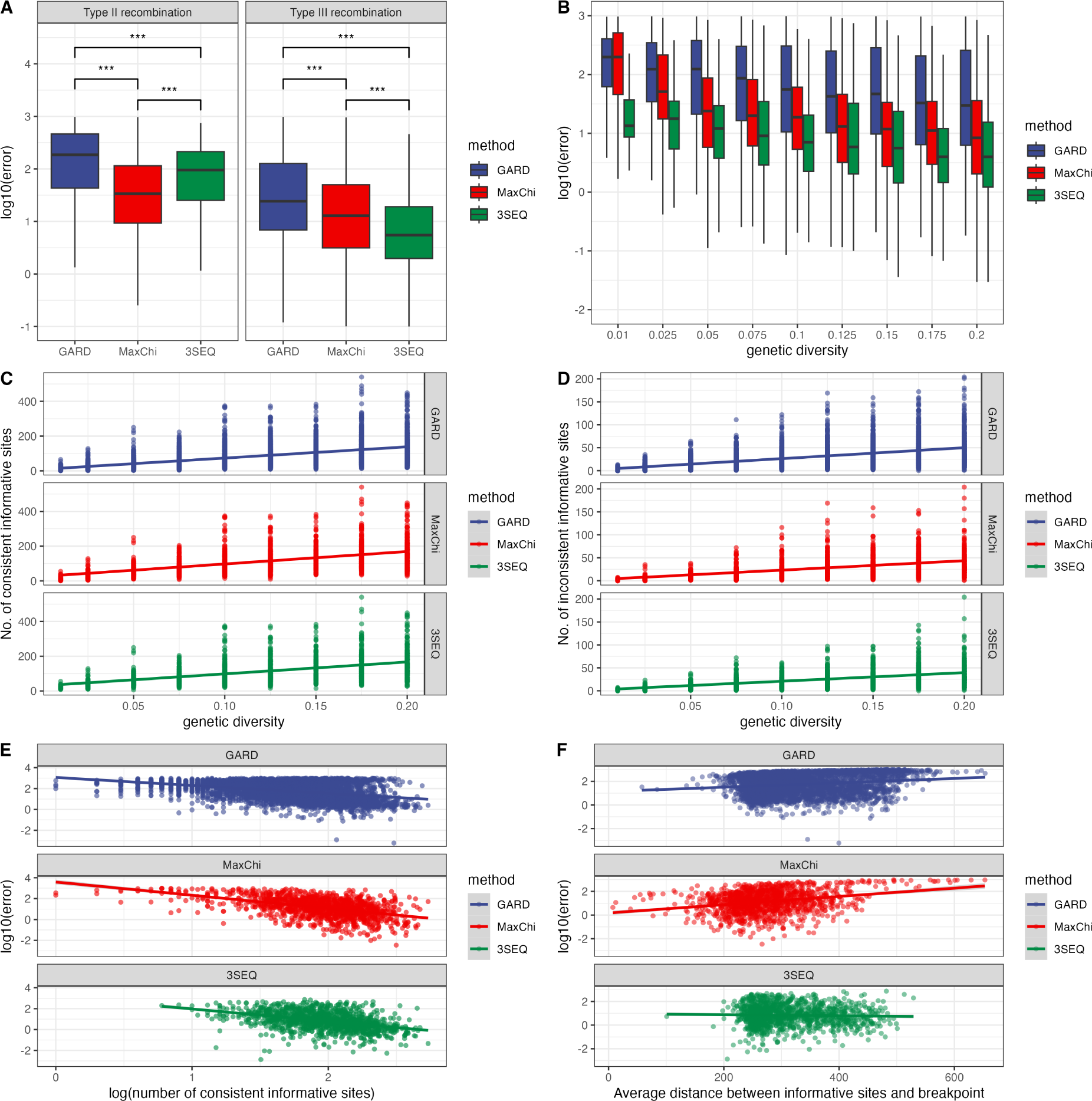
The impact of different genetic features on localization error. (A) Localization error by recombination type. A *** indicates that *P* < 0.001 by ANOVA after Bonferroni correction. (B) Relationship between localization error and genetic diversity. (C) Relationship between genetic diversity and the number of consistent informative sites. (D) Relationship between genetic diversity and the number of inconsistent informative sites. (E) Relationship between localization error and the number of consistent informative sites. (F) Relationship between localization error and average distance between informative sites and the breakpoint.

**Table 4:** Localization accuracy (mean*±*SEM) of three detection methods for multiple breakpoint inference.

|  | Type II | Type III |
| --- | --- | --- |
| GARD | 27.15 ± 1.81 | 23.29 ± 0.63 |
| MaxChi | 11.44 ± 1.98 | 14.76 ± 1.36 |
| 3SEQ | 22.73 ± 1.15 | 20.78 ± 0.66 |

##### Genetic diversity and consistent informative sites

All else being equal, recombination among more genetically divergent sequences should be easier to detect than recombination between genetically similar sequences. In theory then, genetic diversity should closely reflect how much information sequence data contain about recombination. However, as genetic diversity increases the localization error does not decrease as strongly as we might expect, especially for GARD and 3SEQ (Figure 5B).

In order to understand this trend, we further explored the relationship between genetic diversity and the number of informative sites that are either consistent or inconsistent with the allelic state of the parent from which genetic material was inherited in the recombinant child sequence (see Figure 2). As expected, the number of consistent informative sites increases with greater genetic diversity, which should benefit all methods (Figure 5C). However, the number of inconsistent informative sites also increases with genetic diversity (Figure 5D) due to recurrent mutations overriding the correct signal of ancestry. Consistent with these expectations, our PLSR model shows that increasing the number of consistent informative sites decreases localization error, while increasing the number of inconsistent sites increases the localization error. Together these two factors explain up to half of the total variance in performance (Figure 4). Moreover, the regression coefficients for genetic diversity in the PLSR model are non-significant for all three methods when the number of consistent and inconsistent informative sites is included in the model (Table 3). Therefore, while the direct impact of genetic diversity on the localization error is weak, decomposing overall genetic diversity in this way shows that the number consistent and inconsistent sites play a crucial role in determining localization error.

##### Average distance between informative sites and recombination breakpoint

For MaxChi and GARD, the farther away informative sites are from the true recombination breakpoint on average, the higher the localization error (Figure 5F). On the contrary, the distance between informative sites and recombination breakpoint does not significantly affect the error of 3SEQ. The regression coefficients in the PLSR model also match these univariate relationships (Table 3), where the average distance will increase the error for MaxChi and GARD, but have no significant effect on 3SEQ. For MaxChi and GARD it is clear that information about the breakpoint position is maximized when the informative sites are closer to the breakpoint. However, the random walk model employed by 3SEQ appears less sensitive to the distance between informative sites and the breakpoint. As a result, the average distance explains very little variance in the performance of 3SEQ (Figure 4).

#### Precision analysis

We next considered how precisely each method can locate breakpoints based on the probability mass (i.e. site-wise supporting probability) assigned to each site being the breakpoint. For this analysis we only consider type III recombination events. When genetic diversity is relatively low, MaxChi has higher precision than GARD, but as the genetic diversity increases the median precision of both GARD and MaxChi increase and eventually approach one another (Figure 6A). For MaxChi the site-wise probabilities generally increase nearly monotonically towards the breakpoint, but for GARD there are often multiple peaks or local maxima in their distribution (Figure 6B). This could explain why precision varies across data sets much more for GARD than MaxChi. 3SEQ has lower precision than the other two methods, which is expected because 3SEQ assigns all uninformative sites between two informative sites the same site-wise probability, which tends to increase the variance and thus lower precision. For all other methods, precision tends to increase with the number of consistent informative sites, at least when the localization error is low (Figure 6C).

**Figure 6:**
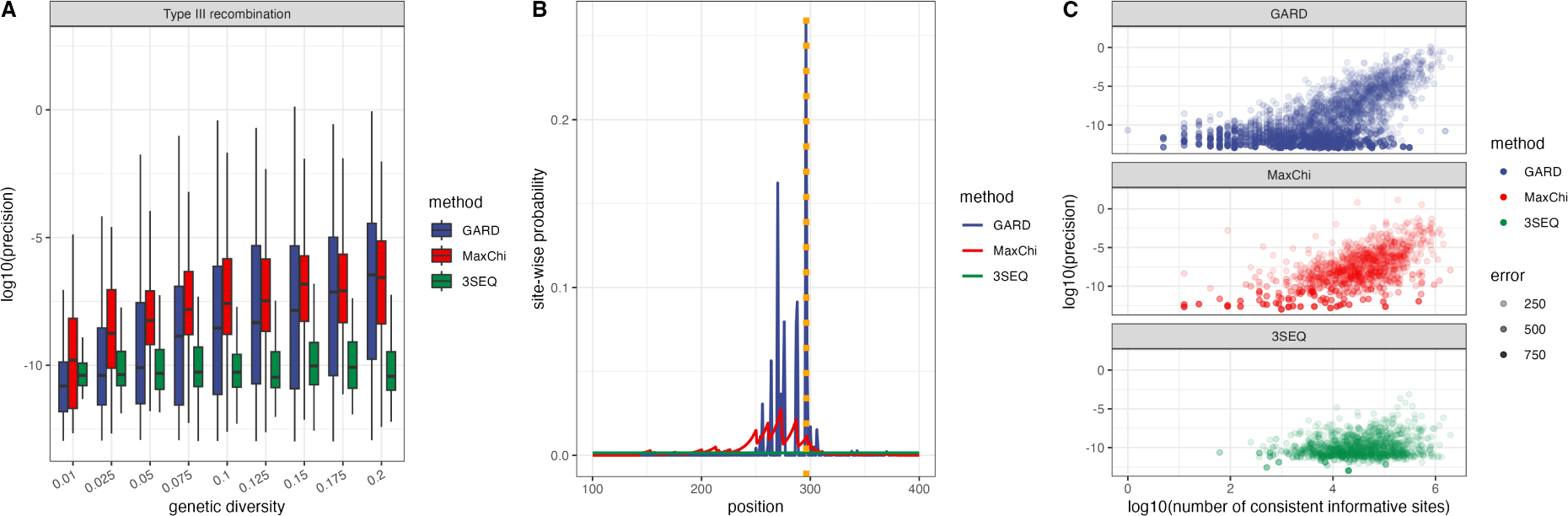
Precision of different detection methods in localizing breakpoints. A. Precision at different levels of genetic diversity; B. An example of how site-wise probabilities are distributed over sites. Here the orange dotted line indicates the true breakpoint location; C. Relationship between the logarithm of precision and the logarithm of the number of consistent informative sites

### Detecting and localizing multiple breakpoints

#### An Overview of the Three Recombination Detection Methods

We compared the power of each method to detect recombination events in simulations with multiple type II and type III recombination events at different levels of genetic diversity (Figure 7). Here, power refers to the ability to detect at least one recombination event. Overall, GARD outperforms the other two methods in terms of power. When genetic diversity is greater than 0.10, GARD always detects a recombination event. When genetic diversity is extremely low, MaxChi and 3SEQ fail to detect any recombination events. With multiple breakpoints, it is not always possible to tell which of the true breakpoints is detected. We therefore quantify the localization error as the distance between each inferred breakpoint and its nearest true breakpoint (see Methods). MaxChi has a slightly lower localization error than 3SEQ and GARD but a larger variance as well (Table 4). We however note that MaxChi’s lower error may result from its limited detection power, such that GARD and 3SEQ have higher errors due to their difficulty localizing breakpoints MaxChi was unable to detect.

**Figure 7:**
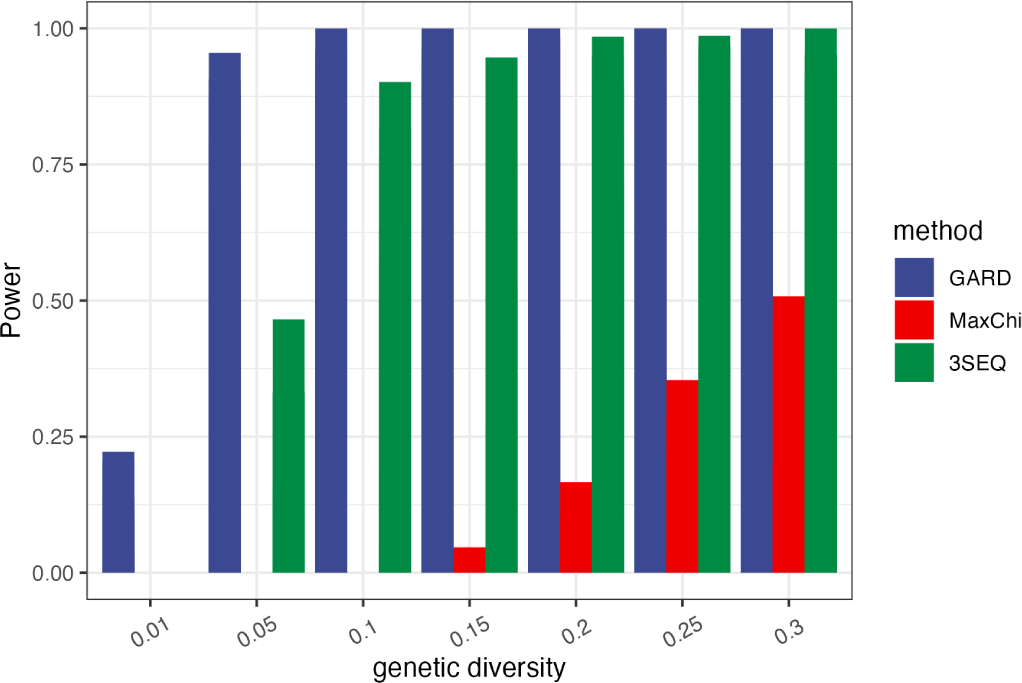
Sensitivity (power) of three methods to detect multiple recombination events

#### Accuracy in localizing breakpoints at different genetic diversity levels

As in our single breakpoint simulations, localization error decreases as genetic diversity increases for 3SEQ and GARD, while the trend for MaxChi is not clear due to its low detection power (Figure 8A). There is no clear relationship between the localization error and the number of breakpoints in an alignment (Figure 8B). Combined with detection power, it seems that 3SEQ performs better than the other two methods as it has a decent detection power as well as a low localization error.

**Figure 8:**
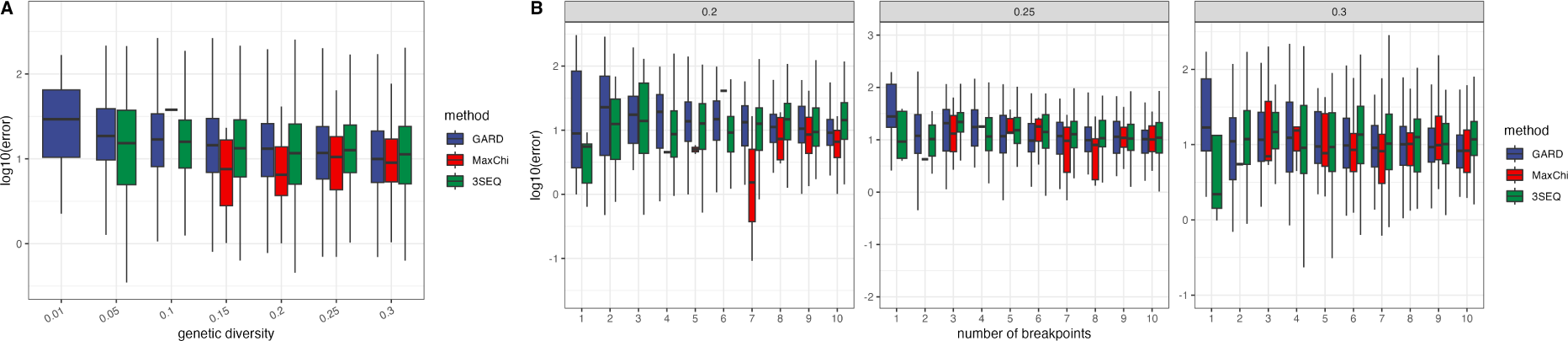
Average localization error in detected breakpoint locations in simulations with multiple break-points. A. Average error across all genetic diversity levels; B. Average error at different levels of genetic diversity and different numbers of breakpoints in the alignment. Only simulations with a genetic diversity greater or equal to 0.20 and type III events are considered as MaxChi cannot detect most recombination breakpoints when the genetic diversity is low.

Based on our single breakpoint results, localization accuracy should decrease with the number of consistent informative sites in the alignment. However, with multiple breakpoints we do not know which true breakpoint is detected, such that defining consistent and inconsistent informative sites is difficult. Hence, we only consider the total number of informative sites. Nevertheless, taking 3SEQ as an example, localization error still decreases with the number of informative sites with multiple breakpoints in the alignment (Figure 9).

**Figure 9:**
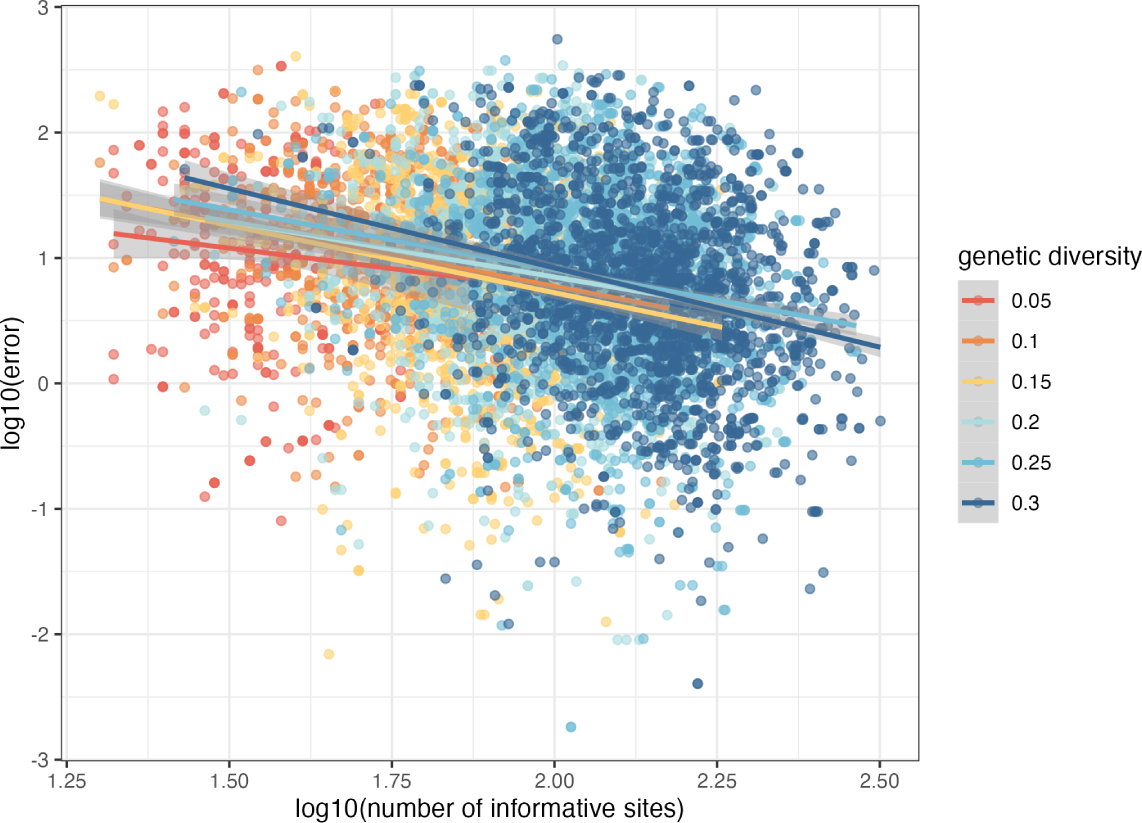
Relationship between the number of informative sites and the breakpoint localization error for 3SEQ in simulations with multiple breakpoints. Linear regression lines are shown for different genetic diversity levels

### Effects of breakpoint localization on phylogenetic reconstruction

The breakpoints identified by recombination detection methods are often used to partition sequence alignments into recombination-free sub-alignments for phylogenetic reconstruction. We therefore wanted to see if using different methods to localize breakpoints would have a measurable impact on the quality of phylogenetic reconstruction. To do this, we ran each detection method on simulated sequence data sets where the underlying true ARG, and thus the true local phylogeny for each region of the genome, was known. Alignments were then cut at the identified recombination breakpoints and maximum likelihood trees were reconstructed from the resulting recombination-free subalignments. To compare performance, we then calculated the Robinson–Foulds distance (RF distance) between each reconstructed tree and the corresponding true local tree at each site in the alignment and then averaged RF distances across all sites. As shown in Figure 10, using the sliced alignment to build the phylogenies can dramatically decrease the average RF distance compared to using the whole alignment which ignores the breakpoints. Generally, the RF distance between the true phylogenies and the phylogenies reconstructed from recombination-free sub-alignments decreases with genetic diversity for all three recombination detection methods before leveling off at higher values of genetic diversity. Comparing the three detection methods, GARD has a slightly better performance than the other two methods but the choice of detection methods has little impact on reconstruction error.

**Figure 10:**
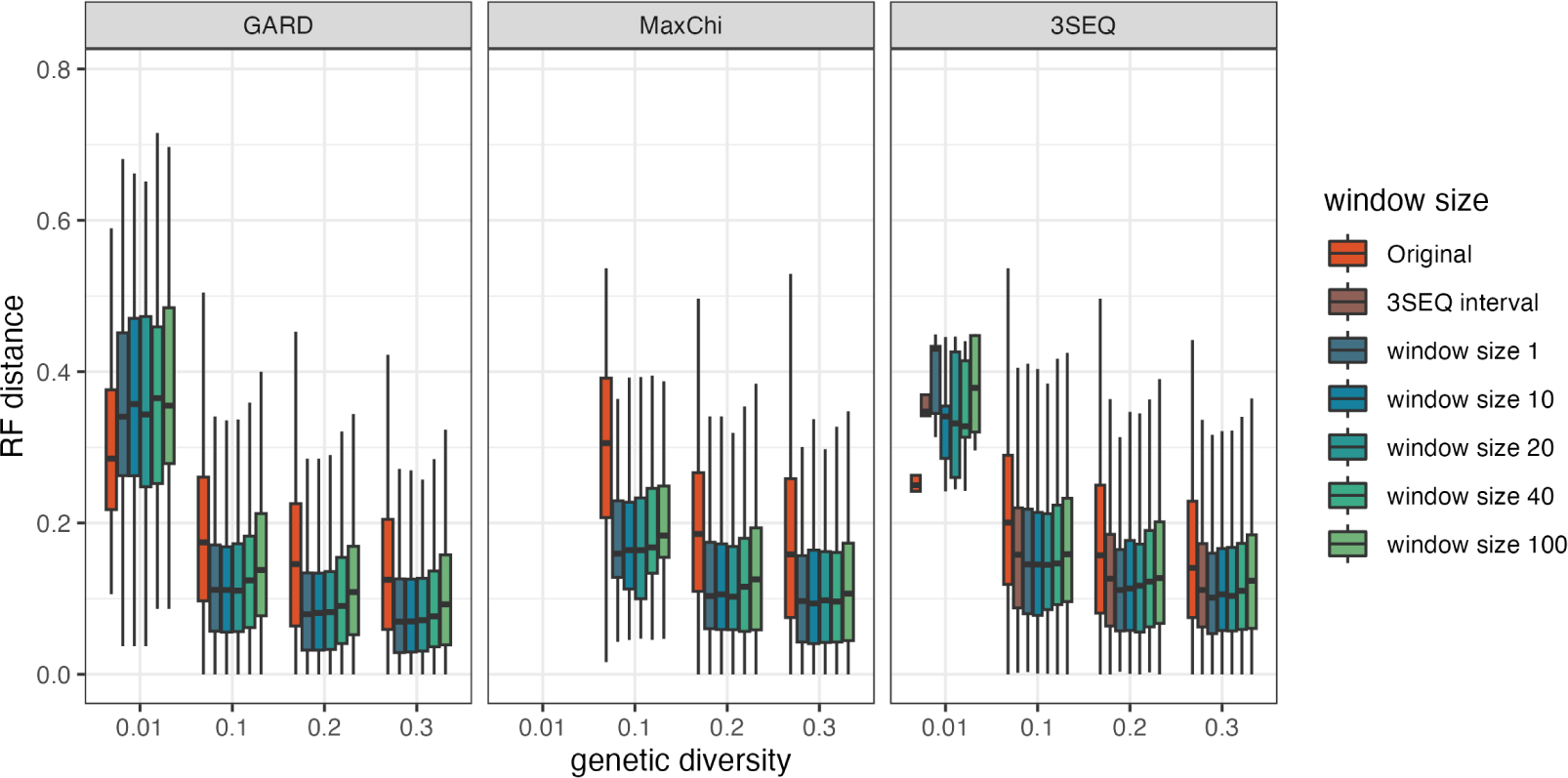
Robinson-Foulds distances between true phylogenetic trees and maximum-likelihood trees reconstructed from subalignments sliced at the recombination breakpoints identified by different methods. Original refers to the full alignment ignoring all breakpoints. Window size refers to the the number of sites neighboring breakpoints that are excluded from the sub-alignments. RF distances between reconstructed and true trees were computed at each site and then averaged across sites. For regions in which a local tree could not be reconstructed due to excluding sites, we use the neighboring ML trees to interpolate between regions when computing RF distances.

We further considered how aggressive or conservative we should be about slicing alignments by excluding sites within a certain distance of identified breakpoints (i.e. the breakpoint window, see Figure S2). Increasing the window size therefore lowers the probability of a true breakpoint being erroneously included in a sub-alignment but at the potential cost of excluding informative sites from the alignment. Overall, RF distances increase slowly as window sizes increase (Figure 10). It is notable that for 3SEQ, using the conservative intervals given by the algorithm to cut the alignment and build the phylogenies is no better than using the mid-point of these intervals. These results suggest that if we are confident about breakpoint locations (Table 4), aggressive strategies may slightly outperform more conservative approaches. We should therefore retain as many sites as possible by slicing alignments as close as possible to identified breakpoints (i.e. choosing a window size of 0).

But overall, the quality of phylogenetic reconstruction depends very little on how we partition the alignment. This can be explained by the fact that there is a trade-off between being aggressive, which can lead to missed breakpoints, and being conservative, which can yield shorter sub-alignments with less informative sites (Supplementary Figure 2). Both a greater number of missed breakpoints and a smaller number of informative sites increase RF distances. Hence, we cannot see a clear difference in RF distances for different window sizes. Accounting for recombination by partitioning alignments at identified breakpoints therefore appears to be much more important than exactly how alignments are partitioned.

In the preceding analysis, the average RF distance across all sites was used to quantify reconstruction accuracy across the entire genome. This performance metric therefore gives more weight to local trees spanning a larger proportion of the genome. On the other hand, if we only care about the the accuracy of individual local trees regardless of what proportion of the genome they span, using a more conservative approach with a larger window size improves reconstruction accuracy for GARD, 3SEQ, or MaxChi (Supplement Figure 3).

## Discussion

In this study, we conducted an in-depth exploration of the performance of three widely used recombination detection methods using simulated ARGs and sequence data. Our investigation revealed that these three methods perform very differently in detecting and localizing recombination breakpoints. For instance, 3SEQ exhibited a tendency toward lower localization errors but also lower precision, while GARD demonstrated robust detection power, albeit with subpar specificity and somewhat unsatisfactory localization accuracy. More generally, we identified several genetic features that can explain this variable performance in breakpoint localization. These features include recombination type and average distance between informative sites and breakpoint. Overall though, the number of (consistent) informative sites was found to be the most important factor determining the accuracy at which both single and multiple breakpoints could be localized. Armed with these insights, we proceeded to evaluate the accuracy of phylogenies reconstructed from sequence alignments partitioned at the breakpoint positions imputed by each method. Notably, we found that the choice of recombination detection method had minimal impact on phylogenetic reconstruction. These findings show that partitioning alignments on identified breakpoints does improve reconstruction accuracy, irrespective of the specific recombination detection method employed.

The statistical models underlying these different methods each consider unique information coming from sequence data as a signal of recombination. The variable performance of these methods can often be explained by features of the genetic data that may amplify or obscure these signals of recombination. For instance, recombination type plays a pivotal role in GARD’s ability to infer breakpoint positions because GARD relies on detecting topological discordance between local phylogenetic trees, which will only be present at Type III recombination events. In contrast, MaxChi considers the clustering of substitutions along pairs of aligned sequences and is therefore much better at detecting Type II recombination events than GARD. However, the accuracy of GARD and MaxChi declines when informative sites are distant from the true breakpoint, as these methods consider the length of local segments on each side of a penitential breakpoint as well during position inference. Conversely, 3SEQ remains less affected by the distance of informative sites. Knowledge of how these genetic features impact performance therefore allows for more informed method selection based on prior information.

Among the genetic features we explored, the number of consistent informative sites emerges as the most influential contributor to localization error, especially when we differentiate between consistent and inconsistent informative sites. As previously reported, detection power increases as genetic diversity and therefore the number of informative sites also increases (Posada and Crandall, 2001), yet breakpoint localization accuracy only increases slowly with increasing genetic diversity. This pattern can be better understood if we view the mutations generating genetic diversity as a “double-edged” sword for detecting recombination. Higher mutation rates create informative sites where the allelic state of a child recombinant sequence correctly matches the allelic state of the parent from which genetic material was inherited, creating a signal of recombination. However, higher mutation rates also create recurrent mutations (homoplastic sites) which obscure patterns of inheritance and thus signals of recombination. Localization accuracy therefore rapidly improves with the number of consistent informative sites but declines with the number of inconsistent informative sites for all three detection methods. While it is not possible to differentiate consistent from inconsistent sites in real sequence data, their significance is undeniable in our simulation study.

Phylogenetic analysis conventionally assumes that the ancestral relationships among samples can be adequately represented by a single phylogenetic tree. However, this assumption often fails when applied to real-world data, leading to inaccurate phylogenetic inference and estimates of molecular clock rates (Schierup and Hein, 2000; Zhu *et al*., 2022). Consequently, recombination detection methods are often used to identify breakpoint locations to generate recombination-free subalignments from which local phylogenetic trees can be reconstructed. Our findings indicate that as long as genetic diversity is not excessively low, there is indeed a significant reduction in phylogenetic error when reconstructing local trees using recombination-free segments (see Figure 10). However, the exact choice of method used to detect breakpoints had little downstream impact on phylogenetic reconstruction accuracy, suggesting that running a single detection method is likely to perform equally as well as running multiple methods. Of the methods we compared, 3SEQ would be our preferred choice as it exhibits robust detection capabilities, high localization accuracy and remains computationally efficient, even for larger datasets. Moreover, we observed remarkably little variation in reconstruction error when different window sizes were used to exclude sites within a certain distance of identified breakpoints. These results suggest that when selecting inferred breakpoints for reconstructing local trees, a more aggressive approach using longer segments may be preferred in order to retain a greater number of informative sites if we want to capture the majority of information of the whole genome (Figure 10). However, being aggressive comes at the cost of potentially including missed breakpoints in the resulting subalignments. The resulting trade-off between subalignment length and the inclusion of missed breakpoints likely explains why the exact partitioning method has little impact on reconstruction accuracy. If we are interested in recovering individual local trees independently of what proportion of the genome they represent, using a more conservative approach to exclude other potential breakpoint sites may be preferred (Supplement Figure 1).

While we only compared three recombination detection methods, the selected methods are representative of the major categories of statistical methods available for both detecting and localizing recombination events. GARD is a classical phylogenetic method, MaxChi falls within the substitution category, and 3SEQ is a triplet-based method. Both MaxChi and 3SEQ are widely used and have been incorporated into population software packages like RDP5 (Martin *et al*., 2020). We however note that the results of these methods may depend on the settings and parameters chosen by the user, especially the significance level chosen as the detection threshold. In practical terms, one can employ a higher significance level (lower p-value) threshold for 3SEQ and MaxChi to increase specificity, potentially leading to different results compared to our current settings. Nonetheless, in our study, we adhered to the default parameters, which are commonly used in practice. Consequently, our results remain representative of the general performance of these detection methods.

Our study also sheds light on how to potentially improve the accuracy and precision of recombination detection methods in the future. For example, one could incorporate information about recombination hot spots or recombination rate variation across sites from empirical studies as an informative prior on the location of breakpoints. More ambitiously, one could estimate the number of consistent informative sites in an alignment based on the genetic diversity and the underlying demographic model. Having prior information about the relative odds of informative sites being generated by true recombination events (consistent sites) rather than recurrent mutations (inconsistent sites) could allow for more accurate and precise inferences which would better reflect our true uncertainty about breakpoint locations.

## Acknowledgements

This research was supported by grant 2019-67021-29932 from the USDA NIFA. DAR was additionally supported by funding from USDA Hatch project 1016556.

## References

Alves, I., Houle, A. A., Hussin, J. G., and Awadalla, P. 2017. The impact of recombination on human mutation load and disease. Philosophical Transactions of the Royal Society B: Biological Sciences, 372(1736): 20160465. recombination impact.

Awadalla, P. 2003. The evolutionary genomics of pathogen recombination. Nature Reviews Genetics, 4(1): 50–60. recombination review.

Baumdicker, F., Bisschop, G., Goldstein, D., Gower, G., Ragsdale, A. P., Tsambos, G., Zhu, S., Eldon, B., Ellerman, E. C., Galloway, J. G., Gladstein, A. L., Gorjanc, G., Guo, B., Jeffery, B., Kretzschumar, W. W., Lohse, K., Matschiner, M., Nelson, D., Pope, N. S., Quinto-Cortés, C. D., Rodrigues, M. F., Saunack, K., Sellinger, T., Thornton, K., Kemenade, H. v., Wohns, A. W., Wong, Y., Gravel, S., Kern, A. D., Koskela, J., Ralph, P. L., and Kelleher, J. 2021. Efficient ancestry and mutation simulation with msprime 1.0. Genetics, 220(3): iyab229.

Bay, R. A. and Bielawski, J. P. 2011. Recombination detection under evolutionary scenarios relevant to functional divergence. Journal of Molecular Evolution, 73(5–6): 273–286. evaluation.

Bell, S. M. and Bedford, T. 2017. Modern-day siv viral diversity generated by extensive recombination and cross-species transmission. PLoS pathogens, 13(7): e1006466.

Bertrand, F. and Maumy, M. 2023. Partial Least Squares Regression for Generalized Linear Models. R package version 1.5.1.

Boni, M. F., Posada, D., and Feldman, M. W. 2007. An Exact Nonparametric Method for Inferring Mosaic Structure in Sequence Triplets. Genetics, 176(2): 1035–1047.

Boni, M. F., Lemey, P., Jiang, X., Lam, T. T.-Y., Perry, B. W., Castoe, T. A., Rambaut, A., and Robertson, D. L. 2020. Evolutionary origins of the sars-cov-2 sarbecovirus lineage responsible for the covid-19 pandemic. Nature microbiology, 5(11): 1408–1417.

Eshelman, L. J. 1991. The CHC adaptive search algorithm: How to have safe search when engaging in nontraditional genetic recombination. In G. J. E. Rawlins, editor, Foundations of Genetic Algorithms, pages 265–283. San Francisco, CA: Morgan Kaufmann.

Haenlein, M. and Kaplan, A. M. 2004. A Beginner’s Guide to Partial Least Squares Analysis. Understanding Statistics, 3(4): 283–297.

Hudson, R. R. 1983. Properties of a neutral allele model with intragenic recombination. Theoretical Population Biology, 23(2): 183–201.

Hudson, R. R. et al. 1990. Gene genealogies and the coalescent process. Oxford surveys in evolutionary biology, 7(1): 44.

Hudson, R. R. and Kaplan, N. L. 1985. Statistical properties of the number of recombination events in the history of a sample of DNA sequences. Genetics, 111(1): 147–164.

Jackson, B., Boni, M. F., Bull, M. J., Colleran, A., Colquhoun, R. M., Darby, A. C., Haldenby, S., Hill, V., Lucaci, A., McCrone, J. T., et al. 2021. Generation and transmission of interlineage recombinants in the sars-cov-2 pandemic. Cell, 184(20): 5179–5188.

Kubatko, L. S. and Degnan, J. H. 2007. Inconsistency of phylogenetic estimates from concatenated data under coalescence. Systematic biology, 56(1): 17–24.

Martin, D. P., Varsani, A., Roumagnac, P., Botha, G., Maslamoney, S., Schwab, T., Kelz, Z., Kumar, V., and Murrell, B. 2020. RDP5: a computer program for analyzing recombination in, and removing signals of recombination from, nucleotide sequence datasets. Virus Evolution, 7(1): veaa087.

Muller, H. J. 1964. The relation of recombination to mutational advance. Mutation Research/Fundamental and Molecular Mechanisms of Mutagenesis, 1(1): 2–9.

Pond, S. L. K. and Frost, S. D. W. 2005. A genetic algorithm approach to detecting lineage-specific variation in selection pressure. Molecular biology and evolution, 22 3: 478–85.

Pond, S. L. K., Posada, D., Gravenor, M. B., Woelk, C. H., and Frost, S. D. W. 2006. Automated Phylogenetic Detection of Recombination Using a Genetic Algorithm. Molecular Biology and Evolution, 23(10): 1891–1901.

Posada, D. 2002. Evaluation of methods for detecting recombination from dna sequences: empirical data. Molecular Biology and Evolution, 19(5): 708–717.

Posada, D. and Crandall, K. A. 2001. Evaluation of methods for detecting recombination from DNA sequences: Computer simulations. Proceedings of the National Academy of Sciences, 98(24): 13757–13762.

Posada, D., Crandall, K. A., and Holmes, E. C. 2002. Recombination in evolutionary genomics. Annual Review of Genetics, 36(1): 75–97.

Rasmussen, M. D., Hubisz, M. J., Gronau, I., and Siepel, A. 2014. Genome-wide inference of ancestral recombination graphs. PLoS Genetics, 10(5): e1004342.

Ratmann, O., Hodcroft, E. B., Pickles, M., Cori, A., Hall, M., Lycett, S., Colijn, C., Dearlove, B., Didelot, X., Frost, S., Md Mukarram Hossain, A. S., Joy, J. B., Kendall, M., Kuhnert, D., Leventhal, G. E., Liang, R., Plazzotta, G., Poon, A. F., Rasmussen, D. A., Stadler, T., Volz, E., Weis, C., Brown, A. J., and Fraser, C. 2016. Phylogenetic Tools for Generalized HIV-1 Epidemics: Findings from the PANGEA-HIV Methods Comparison. Molecular Biology and Evolution, 34(1): 185–203.

Robinson, D. and Foulds, L. 1981. Comparison of phylogenetic trees. Mathematical Biosciences, 53(1-2): 131–147.

Schierup, M. H. and Hein, J. 2000. Consequences of recombination on traditional phylogenetic analysis. Genetics, 156(2): 879–891.

Shikov, A. E., Malovichko, Y. V., Nizhnikov, A. A., and Antonets, K. S. 2022. Current methods for recombination detection in bacteria. International Journal of Molecular Sciences, 23(11): 6257. evaluation.

Smith, J. M. 1992. Analyzing the mosaic structure of genes. Journal of Molecular Evolution, 34(2): 126–129.

Smith, J. M. and Smith, N. H. 1998. Detecting recombination from gene trees. Molecular Biology and Evolution, 15(5): 590–599.

Spielman, S. J. and Wilke, C. O. 2015. Pyvolve: A Flexible Python Module for Simulating Sequences along Phylogenies. PLOS ONE, 10(9): 1–7.

Stamatakis, A. 2014. RAxML version 8: a tool for phylogenetic analysis and post-analysis of large phylogenies. Bioinformatics, 30(9): 1312–1313.

Sukumaran, J. and Holder, M. T. 2010. DendroPy: a Python library for phylogenetic computing. Bioinformatics, 26(12): 1569–1571.

Tamura, T., Ito, J., Uriu, K., Zahradnik, J., Kida, I., Anraku, Y., Nasser, H., Shofa, M., Oda, Y., Lytras, S., Nao, N., Itakura, Y., Deguchi, S., Suzuki, R., Wang, L., Begum, M. M., Kita, S., Yajima, H., Sasaki, J., Sasaki-Tabata, K., Shimizu, R., Tsuda, M., Kosugi, Y., Fujita, S., Pan, L., Sauter, D., Yoshimatsu, K., Suzuki, S., Asakura, H., Nagashima, M., Sadamasu, K., Yoshimura, K., Yamamoto, Y., Nagamoto, T., Schreiber, G., Maenaka, K., Consortium, T. G. t. P. J. G.-J., Ito, H., Misawa, N., Kimura, I., Suganami, M., Chiba, M., Yoshimura, R., Yasuda, K., Iida, K., Ohsumi, N., Strange, A. P., Takahashi, O., Ichihara, K., Shibatani, Y., Nishiuchi, T., Kato, M., Ferdous, Z., Mouri, H., Shishido, K., Sawa, H., Hashimoto, R., Watanabe, Y., Sakamoto, A., Yasuhara, N., Suzuki, T., Kimura, K., Nakajima, Y., Nakagawa, S., Wu, J., Shirakawa, K., Takaori-Kondo, A., Nagata, K., Kazuma, Y., Nomura, R., Horisawa, Y., Tashiro, Y., Kawai, Y., Irie, T., Kawabata, R., Motozono, C., Toyoda, M., Ueno, T., Hashiguchi, T., Ikeda, T., Fukuhara, T., Saito, A., Tanaka, S., Matsuno, K., Takayama, K., and Sato, K. 2023. Virological characteristics of the SARS-CoV-2 XBB variant derived from recombination of two Omicron subvariants. Nature Communications, 14(1): 2800.

Visscher, P. M. and Hill, W. G. 2006. Estimation of recombination rate and detection of recombination hotspots from dense single-nucleotide polymorphism trio data. Genetics, 173(4): 2415–7.

Wiuf, C. and Hein, J. 1999. Recombination as a Point Process along Sequences. Theoretical Population Biology, 55(3): 248–259.

Wiuf, C., Christensen, T., and Hein, J. 2001. A simulation study of the reliability of recombination detection methods. Molecular Biology and Evolution, 18(10): 1929–1939.

Zhu, T., Flouri, T., and Yang, Z. 2022. A simulation study to examine the impact of recombination on phylogenomic inferences under the multispecies coalescent model. Molecular Ecology, 31(10): 2814–2829.

