## Supplemental Material for "Exploring the accuracy and limits of algorithms for localizing recombination breakpoints"

#### Supplementary material

##### 1. Site-wise probability derivation

For 3SEQ, the probability of each site being a recombination breakpoint is calculated as the probability of that site reaching the maximum height in a hypergeometric random walk in the single breakpoint case given $m$ and $n$, the number of informative sites for each parent sequence in sequence triplet. Hence, for each position $\theta_i$, we have

$$P(\theta = \theta_i | m, n) = P(\max \mathbf{H}_{m,n} \leq h_i) = 1 - P(\max \mathbf{H}_{m,n} > h_i) = 1 - \binom{m+n}{n+(h_i+1)} \Big/ \binom{m+n}{n}$$

according to Boni et al. (Barton and Mallows, 1965; Boni *et al.*, 2007).

For MaxChi, the relative site-specific probabilities are derived based on the probability mass function of the hypergeometric distribution and applying the Bayes theorem. Let $L$ be the length of the genome, $\theta$ be the inferred position and $K$ be the total number of base pairs with different states. For a specific position $\theta_i \in (0, L)$, let $K_{\theta_i}$ be the number of base pairs with different states on the left side of position $\theta_i$. Hence we have the posterior distribution of $\theta$ given the sequences and K,

$$\begin{aligned} P(\theta = \theta_i | Sequences, K) &= \frac{Likelihood(Sequences | \theta = \theta_i, K)\pi(\theta)}{m(Sequences)} \\ &\propto Likelihood(Sequences | \theta = \theta_i, K) \\ &= \frac{1}{\binom{\theta_i}{K_{\theta_i}}\binom{L-\theta_i}{K-K_{\theta_i}}} \propto \frac{1}{\frac{\binom{\theta_i}{K_{\theta_i}}\binom{L-\theta_i}{K-K_{\theta_i}}}{\binom{L}{K}}} \\ &= \frac{1}{P(X = K_{\theta_i})}. \end{aligned}$$

where $X \sim \text{Hyper}(K, \theta_i, L)$ and $\theta \sim Unif(0, L)$ based on Smith et al.'s model assumption (Smith, 1992). We used the p-value from the $\chi^2$ test of the $2 \times 2$ contingency table to approximate $P(X = K_{\theta_0})$.

For GARD, the site-specific probabilities are given in the program output.

#### 2. Supplementary Figures

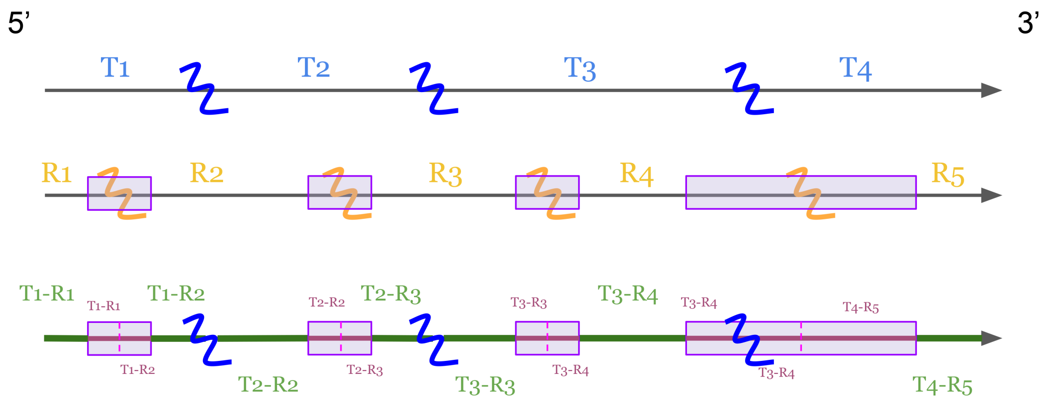


Supplementary Figure 1: Illustration of how Robinson-Foulds distances are computed between true and reconstructed trees. The first alignment is partitioned by true breakpoints, generating three true local trees $(T_1, T_2,$ and $T_3)$. The second alignment is partitioned by inferred breakpoints $(R_1, R_2, R_3$ and $R_4)$. The third alignment shows how RF distance is calculated between true local trees and reconstructed trees.

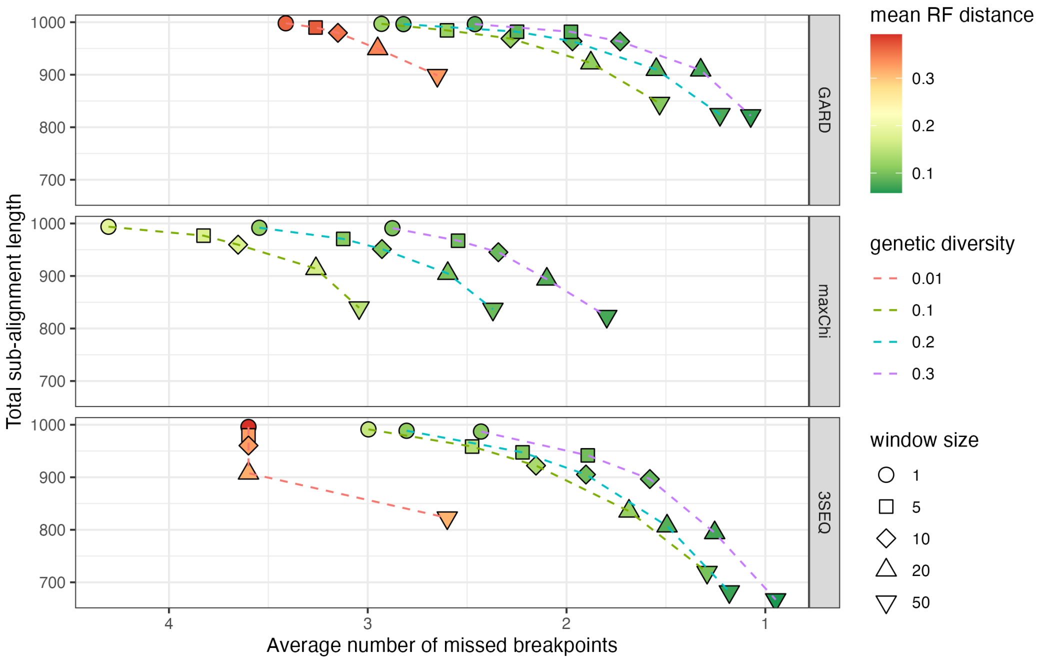


Supplementary Figure 2: The scatter plot between RF distance averaging over all simulations and the average number of missed breakpoints in recombination-free sub-alignments

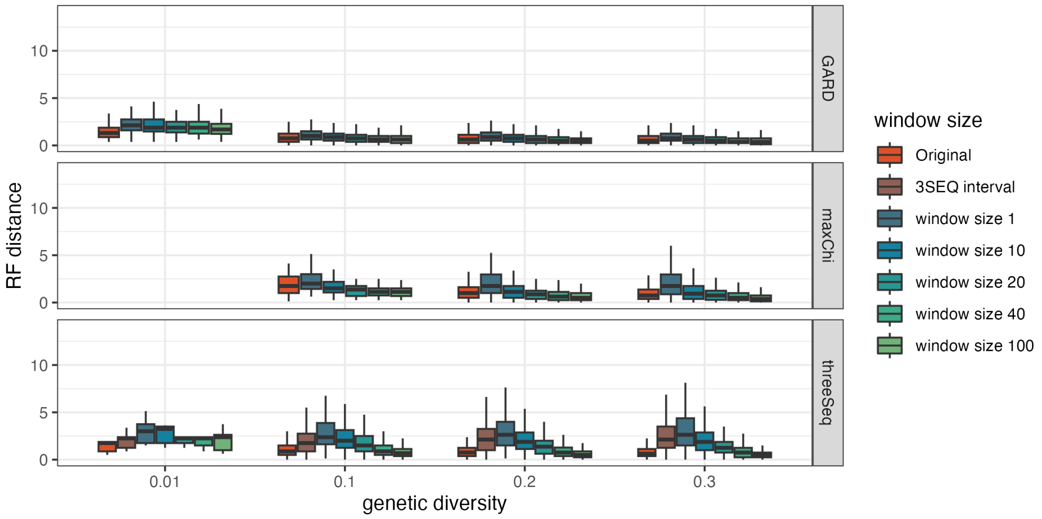


Supplementary Figure 3: The boxplot of total RF distance between the true phylogenies and split alignments at different genetic diversity levels for different window sizes.

### References


Barton, D. E. and Mallows, C. L. 1965. Some Aspects of the Random Sequence. *The Annals of Mathematical Statistics*, 36(1): 236 – 260.

Boni, M. F., Posada, D., and Feldman, M. W. 2007. An Exact Nonparametric Method for Inferring Mosaic Structure in Sequence Triplets. *Genetics*, 176(2): 1035–1047.

Smith, J. M. 1992. Analyzing the mosaic structure of genes. *Journal of Molecular Evolution*, 34(2): 126–129.
